# Is environmental enrichment beneficial for laboratory animals? A systematic review of studies in zebrafish

**DOI:** 10.1101/2023.02.02.526810

**Authors:** Matheus Gallas-Lopes, Radharani Benvenutti, Nayne I. Z. Donzelli, Matheus Marcon

**Author notes:** Correspondence: Matheus Marcon, Ph.D., Departamento de Bioquímica, Farmacologia e Fisiologia, Instituto de Ciências Biológicas e Naturais, Universidade Federal do Triângulo Mineiro (UFTM), Praça Manoel Terra, 330, Uberaba, MG, 38025-015, Brazil.

## Abstract

Environmental enrichment (EE) consists of a series of interventions that are carried out in the home environment to provide greater exposure to sensory stimuli with the objective of mimicking the natural habitat for the animals housed in the laboratory, offering a more complex environment like those found in nature. Some studies have shown the positive effects of EE on zebrafish housed in a laboratory environment. However, this evidence is still very recent and accompanied by contradictory results. Furthermore, there is great variability in the protocols applied, and in the conditions of the tests, tanks, and materials used for creating an EE environment. This substantial variability can bring many uncertainties to the development of future studies and hinder the reproducibility and replicability of research. In this context, the main objective of this study was to carry out a systematic review of the literature aiming to provide an overview of the EE protocols used in zebrafish. We conducted a systematic review following PRISMA recommendations. The literature search was performed in PubMed, Scopus, and Web of Science databases and the studies were selected based on inclusion/exclusion criteria. We performed data extraction and risk of bias analysis of the studies included. A total of 901 articles were identified in the databases and 27 of these studies were included in this review. Among these studies, the effect of EE was evaluated as two different proposals. (1) to improve animal welfare and (2) as an intervention for the prevention of some disorders. Although the zebrafish EE protocols presented a series of experimental differences, the results showed that the benefits of the EE for zebrafish were robust. According to the results described here, the use of EE in the zebrafish home tank provides better welfare and may reduce sources of bias in scientific experiments, such as high-stress levels and fighting events.

## 1 INTRODUCTION

For decades, mammalian species (especially laboratory rodents) have been used as models for the study of human brain disorders (Ellenbroek & Youn, 2016). However, recent translational research in this field is witnessing a growing interest in new models such as flies, roundworms, and fish (Dawson et al., 2018). Especially, zebrafish (*Danio rerio*, Hamilton 1822), a freshwater teleost fish from Southeast Asia commonly found in shallow lakes and rice paddies (Spence et al., 2008), is increasingly attracting the attention of researchers due to its numerous advantages. Mainly, maintenance for this species is more practical when compared with rodents and, besides that, its small size, external development, and genetic similarity to humans (approximately 70%) have favored and contributed to the growing use of this model organism in neuroscience research (Kalueff et al., 2014; Stewart et al., 2014). However, the increasing use of this species in the laboratory environment also raised many concerns about how to maintain its well-being in research facilities (Lee et al., 2022).

Historically, there is a growing concern about animal welfare used in scientific research (Sert et al., 2020), and enriching the home environment has been suggested as a strategy to improve animal welfare (Kempermann, 2019). Environmental enrichment (EE) consists of a series of interventions that are carried out in the home environment to provide greater exposure to sensory stimuli (e.g., visual, motor, olfactory, auditory, and gustatory). Thus, EE aims to mimic the natural habitat for the animals housed in the laboratory, offering a more complex environment that provides physical and mental challenges like those found in nature, increasing individual-environment interaction (Nithianantharajah & Hannan, 2006).

Some studies have shown the positive effects of EE on zebrafish housed in a laboratory environment. For example, EE-housed zebrafish showed an increase in the proliferation of telencephalon cells (von Krogh et al., 2010), and an increase in the size of the brain (DePasquale et al., 2016), corroborating with the results found in rodents, which showed that the central nervous system is more developed when animals were housed in EE condition (Hirase & Shinohara, 2014; Mahati et al., 2016; Zhang et al., 2018). However, although there is some evidence showing the positive effects of EE, there are also contradictory results and there is great variability in the protocols applied, and in the conditions of the tests, tanks, and materials used for creating an EE environment.

This substantial variability can bring many uncertainties to the development of future studies and hinder the reproducibility and replicability of research. In this context, the main objective of this manuscript was to carry out a systematic review of the literature aiming to provide an overview of the EE protocols used in zebrafish, presenting, and discussing: the outcomes evaluated; the results found; the applicability of this intervention in the zebrafish research laboratory routine; and the current and future challenges of the research in this area.

## 2 METHODS

### 2.1 Search strategy

We conducted a systematic review following PRISMA recommendations (Page et al., 2021). On January 15th, 2022, we performed the literature search in PubMed, Scopus, and Web of Science using keywords that contemplate our research topic for the intervention (environmental enrichment) and the desired population (zebrafish), thus we applied the following search term: (‘environmental enrichment’ OR ‘enriched environment’) AND zebrafish. The search was limited from January 1st, 2010, to December 31st, 2021. We also verified reference lists of included studies in this review to detect potentially relevant manuscripts and included the studies published in the time interval between the literature search and the preparation of this manuscript. Database search files are available at the study repository in Open Science Framework (https://osf.io/2gdey/).

### 2.2 Eligibility screening

After searching the studies in the three databases, the selection of studies included in this systematic review was performed by two independent researchers, and the disagreements were discussed with a third reviewer. Initial screening was performed by reading titles/abstracts to identify and exclude duplicates. After excluding duplicates, studies were selected based on inclusion and exclusion criteria, initially after reading the title/abstract and, when necessary, after reading the full text. Studies were included in this review if they satisfied the following criteria: (1) experimental studies that evaluated the effects of EE on zebrafish; (2) studies that considered EE as a complex intervention and used various EE objects; (3) studies reported in English. The exclusion criteria were: (1) studies with EE and no use of zebrafish as an animal model; (2) studies that did not use any animal; (3) studies without adequate control groups; (4) review articles, retracted articles, book chapters, letters, and conference abstracts.

### 2.3 Data extraction

Two investigators worked independently to complete the data extraction, and disagreements were discussed with a third reviewer. All data extraction was performed from the full text or figures. The following information was collected: (1) general data: title, the surname of the first author, publication year, the test used, outcome evaluated, other interventions used, and main results; (2) EE protocols: type of EE intervention (welfare or prevention), materials used, exposure time, use or no of rotation of EE objects; and the dimensions of the EE tank. (3) Experimental details: feeding diet, zebrafish strain used, age of animals, sex of animals, habituation period, water parameters, and light/dark cycle. Co-authorship networks were constructed using VOSviewer software version 1.6.18 (https://www.vosviewer.com) (van Eck & Waltman, 2007, 2010).

### 2.4 Risk of bias

Risk of bias assessment was conducted for each of the studies by two independent authors to evaluate the methodological and reporting quality of the papers included in the review. Disagreements were resolved by discussion between investigators. The analysis was conducted based on the SYRCLE’s risk of bias tool for animal studies (Hooijmans et al., 2014) with adaptations. The following items were evaluated for methodological quality: (1) adequate randomization process for the allocation of animals to experimental groups; (2) comparable baseline characteristics of the animals in each group; (3) blinded outcome assessment by investigators; (4) incomplete or missing data; (5) selective outcome reporting. The reporting quality was evaluated based on the following items: (6) a report of sample size estimation methodology; (7) a report of outlier exclusion criteria. Items 1 to 5 were evaluated as presenting a low, high, or unclear, risk of bias. Items 6 and 7 assessing reporting quality of included studies were scored only as presenting a low or high risk of bias.

## 3 RESULTS

### 3.1 Studies Selected

We have identified a total of 901 articles in the selected databases: PubMed (328), Scopus (197), and Web of Science (376). 426 duplicate records were excluded, 442 records were excluded after title/abstract reading, and 9 studies were excluded after full-text analysis. A total of 24 studies were selected, and we also included 3 more manuscripts published in the time interval between the literature search and the preparation of this manuscript (January 2022 to October 2022), totalizing 27 studies as demonstrated in figure 1.

**Figure 1.**
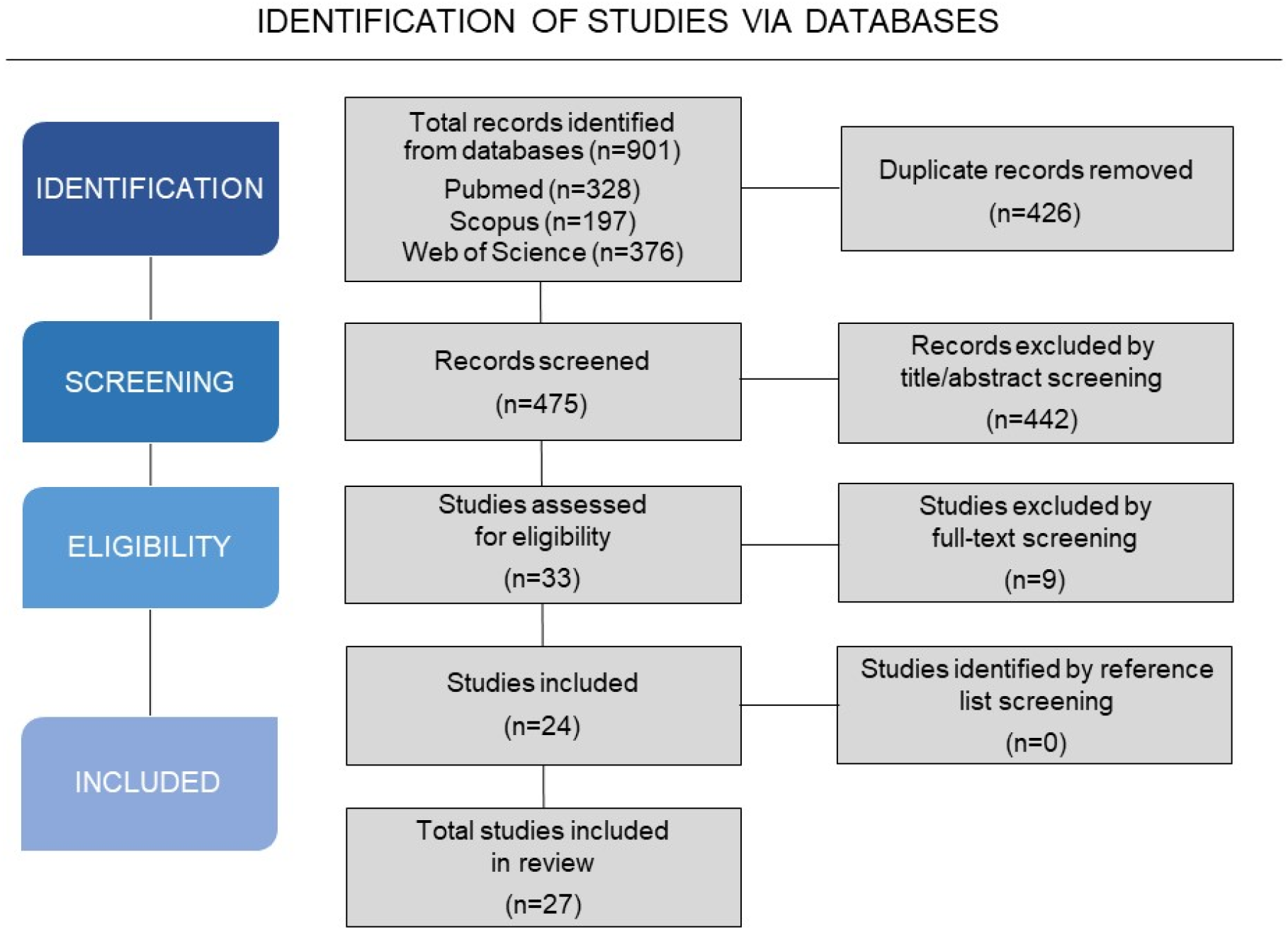
Flowchart diagram of the screening of studies and selection process.

Of the 27 studies included in this review, the year with the lowest number of publications was 2010, with only 1 publication this year. However, as the years progressed, there was an increase in the number of publications on this topic (EE and zebrafish), reaching the highest number of publications per year in 2018 and 2021, with 4 publications each year (Figure 2A). Most of these studies evaluated the potential of EE as a strategy to improve animal welfare, while other studies evaluated the potential of EE as a prevention strategy for the development of some behavior and neurochemical disorders (Figure 2B). In addition, most of these studies aimed to identify the effects of EE on zebrafish behavior, followed by biochemical, developmental, reproductive, molecular, and survival analyses (Figure 2C).

**Figure 2.**
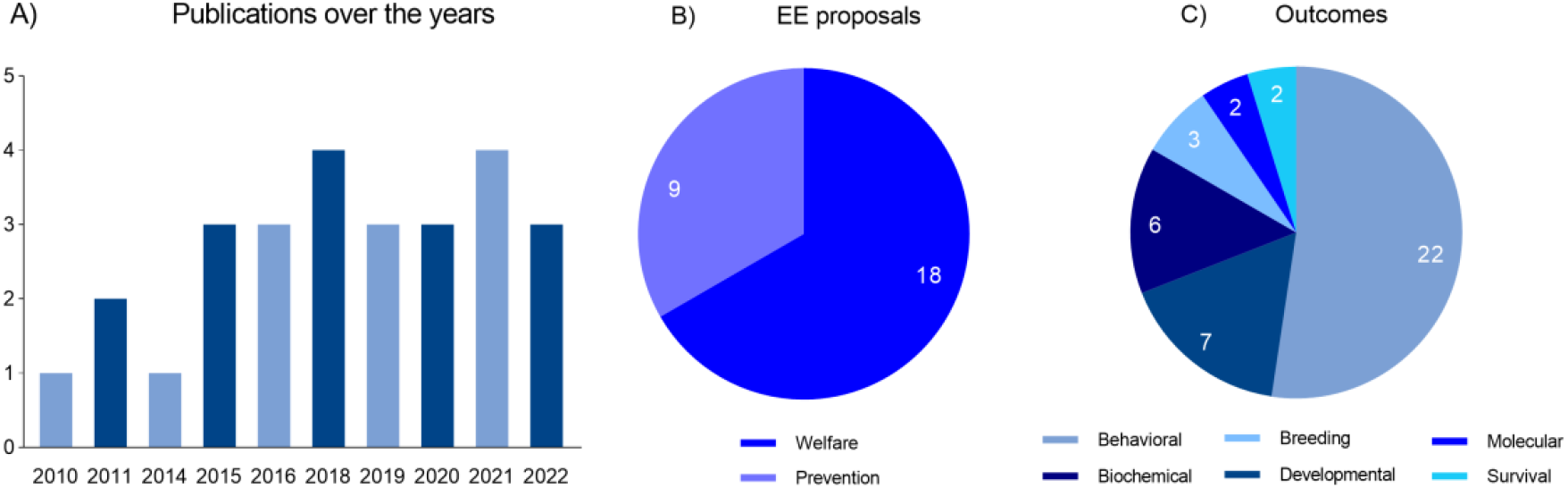
Overview of the publications in peer-reviewed scientific journals that report the use of EE in zebrafish. A) Number of publications over the years. B) Types of EE proposals. C) Evaluated outcomes.

Co-authorship network analysis identified 20 clusters of researchers implementing EE in their labs across the globe based on the studies included in this review (Figure 3). An interactive version of the co-authorship network is available at https://tinyurl.com/2jdo264z.

**Figure 3.**
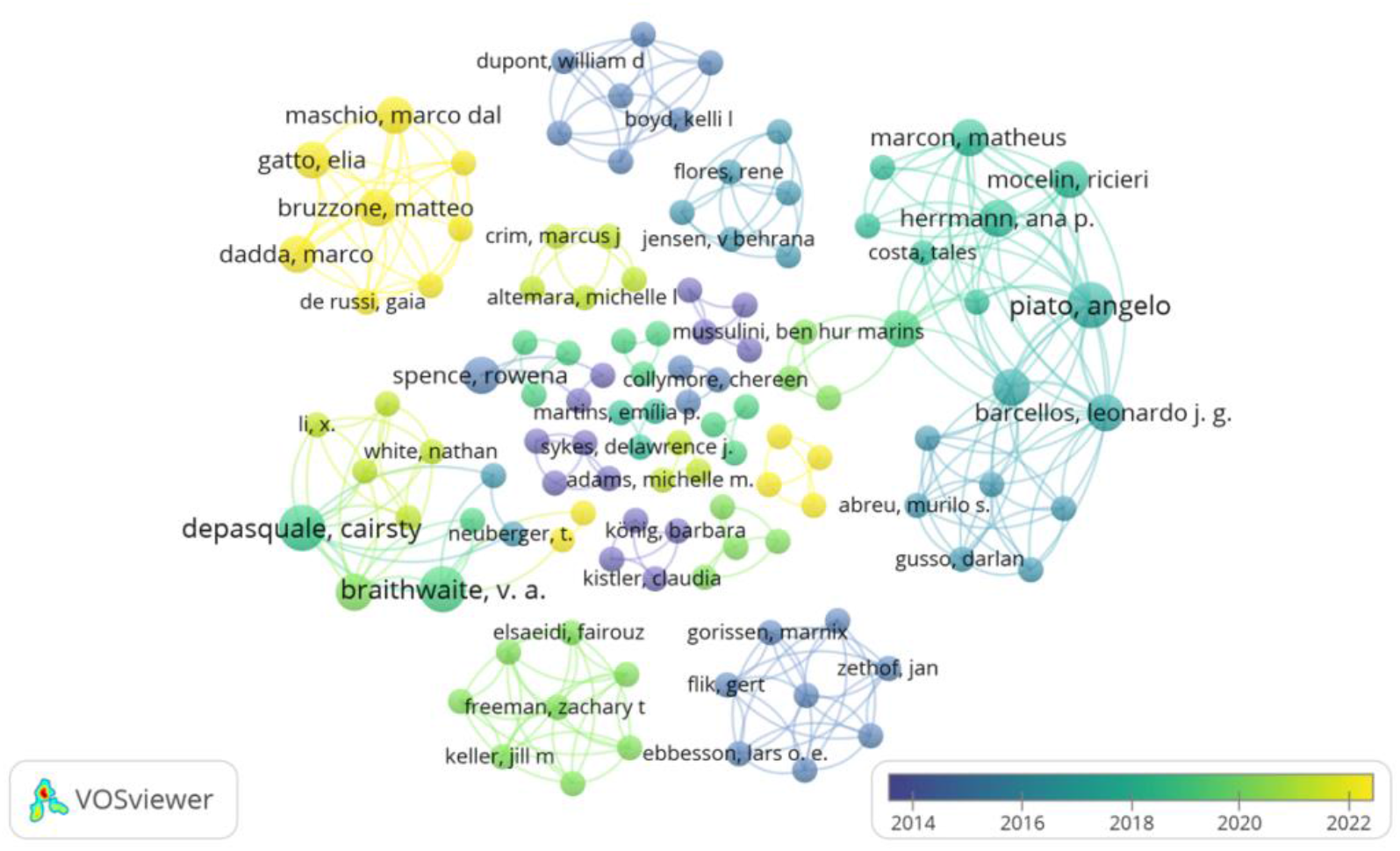
Co-authorship network analysis of researchers that authored studies implementing environmental enrichment (EE) in research with zebrafish. Authors are color-coded from violet (older studies) to yellow (more recent studies) indicating the average publication year of the studies published by each researcher. The size of the circles represents the number of studies published by each author. The distance between the two circles indicates the correlations between researchers.

### 3.2 EE and zebrafish

Historically, it was in the 1920s that the first reports on the importance of EE emerged through a manuscript that tells the story of primates bred in a property in Havana (Yerkes, 1925). The application of EE has also long been reported by zoos and by other animal caretakers to improve animal welfare (Hancocks, 1980; Hutchins et al., 1984). Although the use of EE is remote, it was not long ago that the term “Environmental enrichment (EE)” was recognized and has been discussed as a strategy to improve laboratory animal welfare (Beaver, 1989). Recently, in addition to many studies investigating the potential of environmental enrichment as a strategy to improve animal welfare, many studies have also emerged evaluating and discussing the potential of this intervention as a non-pharmacological therapeutic approach for the treatment of other disorders, for example, mood disorders (Hegde et al., 2020; Huang et al., 2021; Jha et al., 2011; Seong et al., 2018), and neurodegenerative diseases (Griñán-Ferré et al., 2018; Jankowsky et al., 2005; Seo et al., 2020; Wassouf et al., 2018). In zebrafish, until this date, the studies in the literature have evaluated the effects of EE as two different types of proposals: (1) to improve zebrafish welfare and (2) for prevention of behavior, biochemical, molecular, development, and breeding disorders.

#### 3.2.1 EE as a welfare strategy

As preclinical research practices evolved throughout the years, special attention has been given by researchers and institutions to maximizing the welfare of animals used in experiments. As mentioned before, in a husbandry and housing lab facility, EE is applied to diminish the negative impacts of this circumstance on animal welfare and mimic some conditions of the natural habitat of the species in question. Research using zebrafish is no exception, and various studies in this review assessed the impacts of EE as a welfare strategy to improve the quality of life and development of this model organism in experimental conditions. Table 1 summarizes the articles reviewed in this section.

**Table 1.** Studies included in this review that evaluated the implementation of EE to improve zebrafish welfare.

| OUTCOME | INTERVENTIONS | ASSAY | MAIN RESULTS | STUDY ID |
| --- | --- | --- | --- | --- |
| **Behavioral** (locomotor)<br>**Biochemical** (cortisol) | No | Locomotor activity,<br>Whole-body cortisol, Body | EE ↑ the number of PCNA in the telencephalon and ↑ slightly the cortisol levels. | von Krogh<br>*et al.* 2010 |
| **Molecular** (*pcna*)<br>**Developmental** (growth and cell proliferation) |  | length, and weight,<br>Immunohistochemistry. |  |  |
| **Behavioral** (preference) | No | Place preference. | Zebrafish preferred an EE place over a barren place. | Kistler *et al.* 2011 |
| **Behavioral** (learning)<br>**Development** (growth) | No | Maze associative learning,<br>Body length. | Zebrafish raised in EE showed ↑ learning and were smaller than those raised in barren conditions. | Spence *et al.* 2011 |
| **Behavioral** (preference) | Different housing conditions (pairs or groups) | Place preference. | Dominant zebrafish housed in pairs and the zebrafish housed in groups preferred the EE place. Gravel substrate and gravel images were the EE type preferred. | Schroeder *et al.* 2014 |
| **Biochemical** (cortisol)<br>**Survival** | No | Whole-body cortisol,<br>Death count. | EE prevented death in pair-housed zebrafish; EE for 5 days ↑ cortisol levels and EE for 10 days prevented the ↑ of cortisol levels in pair-housed zebrafish. | Keck *et al.* 2015 |
| **Breeding** (fertility and fecundity) | No | Egg and fry count. | The egg number and the fry number were ↑ for the zebrafish breeding pair placed in EE. | Wafer *et al.* 2016 |
| **Behavioral** (social) | No | Social interaction. | EE ↑ the zebrafish social interaction. | Sykes *et al.* 2018 |
| **Behavioral** (preference) | No | Place preference. | Zebrafish preferred the EE + water flow zone. | DePasquale *et al.* 2019 |
| **Behavioral** (preference) | No | Place preference. | Zebrafish showed no preference for EE type (shelter and artificial plants). | Jones *et al.* 2019 |
| **Behavioral** (anxiety)<br>**Developmental** (growth)<br>**Survival** | No | Novel tank,<br>Body length and mass,<br>Count of live juveniles. | EE ↓ the adult zebrafish anxiety-like behavior, ↑ the female body condition, and ↑ the larvae survival. | Lee *et al.* 2019 |
| **Behavioral** (aggressiveness, boldness, locomotor activity)<br>**Breeding** (fertility and fecundity)<br>**Developmental** (growth) | No | Mirror test,<br>Open field,<br>Body length,<br>Egg fertilized count. | EE ↑ the aggressiveness and ↓ the body length of zebrafish. | Woodward *et al.* 2019 |
| **Behavioral** (anxiety, locomotion, preference) | No | Novel tank,<br>Black/white,<br>Social preference. | EE for 7 days ↓ the total distance traveled, and EE for 14 days did not alter behavior. EE for 14 days with changing the EE stimulus at day 8 ↓ total distance traveled. | dos Santos *et al.* 2020 |
| **Behavioral** (preference) | No | Place preference. | Zebrafish preferred the EE type: mirrored paper; marbles; tulle; and PVC pipe. | Krueger *et al.* 2020 |
| **Behavioral** (preference) | No | Place preference. | Zebrafish preferred an EE place over a barren place. | Tan *et al.* 2020 |
| **Behavioral** (cognition) | No | Novel object recognition. | EE ↑ the object recognition memory. | DePasquale *et al.* 2021 |
| **Behavioral** (anxiety) | No | Novel object exploration,<br>Novel odor exploration. | EE ↓ anxiety-like behavior in 14 and 21 dpf zebrafish larvae. | Gatto, Dadda, *et al.* 2022 |
| **Behavioral** (memory) | No | Novel object recognition. | Zebrafish raised in EE equally explored familiar and novel stimuli (memory impairment). | Gatto, Bruzzone, *et al.* 2022 |
| **Behavioral** (preference) | No | Place preference,<br>Choice test. | Zebrafish showed EE place preference and this preference was maintained in a choice test with a psychological stressor. | Maia *et al.* 2022 |

EE: environmental enrichment; DPF: days post-fertilization; PCNA: proliferating cell nuclear antigen; ↑: increased or higher as compared to the control group; ↓: decreased or lower as compared to the control group.

When given the choice between enriched or barren environments, researchers have shown that zebrafish prefer complex habitats (DePasquale et al., 2019; Kistler et al., 2011; Tan et al., 2020), spending more time in EE zones with combined forms of enrichment, even after exposure to psychological stressors (Maia et al., 2022), showing the benefits of introducing EE as a standard practice to account for the well-being of the animals. Preference for complex environments might be modulated by dominance status (with subordinate fish occupying barren areas to avoid contact with conspecifics) and sex, as male and female fish show a preference for different enrichment options (Schroeder et al., 2014). Jones and collaborators (Jones et al., 2019), on the other hand, reported that zebrafish avoided both shaded areas and plants introduced to serve as a shelter, spending more time in the barren zone of the apparatus. Enrichment options go way beyond artificial plants and gravel, with animals interacting strongly with various options of materials such as marbles, PVC pipes, tulle, and, especially, items that resemble or promote social behaviors such as mirrored paper (Krueger et al., 2020).

Stress is characterized by a cascade of physiological events induced by a perception of a threat. In fish, cortisol is used as a stress response biomarker, and its levels increase during exposure to acute or chronic stress situations (Ramsay et al., 2009). Von Krogh and collaborators (von Krogh et al., 2010) showed that socially isolated male zebrafish housed in aquariums with EE for 7 days showed higher cortisol levels when compared to fish housed in barren environments. However, this increase in whole-body cortisol seen was below those seen in experimentally stressed fish. Animals housed in EE also presented an increase in the number of proliferating cell nuclear antigen (PCNA) positive cells in the telencephalon, potentially indicating a change in proliferative activity in the brain. Whilst evaluating pair-housed zebrafish, Keck and collaborators (Keck et al., 2015) also reported an increase in cortisol levels by day 5 of exposure to EE when compared to single-housed animals and pair-housed animals without enrichment. This increase in cortisol levels significantly decreased by day 10, with levels lower than the other groups. Housing enrichment also prevented deaths and wounds in pair-housed zebrafish (Keck et al., 2015).

Housing conditions have also been shown to influence the cognitive performance and social behavior of zebrafish. Animals raised in a complex environment were able to learn faster to locate food rewards in a five-chambered maze when compared to fish reared in a barren environment (Spence et al., 2011). In the novel object recognition test, where the exploration time of a novel vs. a familiar object is quantified (Antunes & Biala, 2012), some researchers reported that EE enhanced the capacity of adult fish to discriminate between novel and familiar objects (DePasquale et al., 2021) while in larvae, the complex environment reduced the expected neophobic response to an unfamiliar object (Gatto, Bruzzone, et al., 2022). As zebrafish are highly social animals, forming intricate hierarchies both in the wild and in lab facilities, alterations in parameters related to these behaviors can be useful tools to study human disorders (Gerlai, 2014; Suriyampola et al., 2016). Enrichment protocols affect social cohesion as fish raised in EE show increased social interaction response when compared to animals raised in barren tanks, emphasizing the importance of the complexity of the environment on zebrafish behavior (Sykes et al., 2018).

In the novel tank test, zebrafish are exposed to an unfamiliar arena with a water column high enough to evaluate geotaxis. Zebrafish show an innate preference for the bottom zone of the apparatus and the time spent in this zone is used as an indicator of anxiety, while time spent and entries in the upper zone are frequently used to evidence the effects of anxiolytic interventions (Levin et al., 2007). In this test, EE reduced the exploration of the bottom part of the arena, being able to decrease anxiety-like behavior (Lee et al., 2019). Other experiments show that EE has different effects depending on exposure time, with protocols of 7 days reducing the total distance traveled by fish in the novel tank while 14 days of exposure did not alter behavior (dos Santos et al., 2020). When exposed to a novel object, EE larval zebrafish also showed a decrease in the avoidance of the unfamiliar object, which translates to a reduction in anxiety-like behaviors (Gatto, Dadda, et al., 2022). In other commonly used behavioral tests adapted to zebrafish, such as the mirror-induced aggressiveness test, EE-raised fish were significantly more aggressive, showing an increase in the mean number of aggressive attacks they made toward their image in the mirror (Woodward et al., 2019).

Differences in the complexity of the habitat are also able to differently regulate morpho and physiological parameters. Some researchers report that housing in EE led to smaller zebrafish when compared to fish raised in barren environments (Spence et al., 2011; Woodward et al., 2019) as others show an enhancement only of female body condition scores and an increase of larval survival (Lee et al., 2019). Enrichment improves the fecundity and fertility of the animals leading to higher numbers of eggs and fry (Wafer et al., 2016).

The comprehensive literature search conducted in this study stresses the importance of EE as a standard practice for zebrafish housing facilities to maximize the well-being of the animals. The fish significantly prefer environments with different levels of complexity, which also translates into an enhancement in the performance of behavioral tasks evaluating cognitive function and anxiety-like states while also improving body condition, fertility, and survival chances, all pointing to the benefits of this intervention.

#### 3.2.2 EE as a prevention strategy

The zebrafish is an organism model widely used in the neuroscience and pharmacology field as a tool to study neurological conditions and develop new treatments. Aversive and harmful stimuli are known to disrupt the normal behavior of zebrafish, mimicking relevant aspects of psychiatric diseases such as depression and anxiety disorders (Kalueff et al., 2014). Thereby, several studies reviewed here evaluated the potential of EE as an intervention to prevent behavioral, biochemical, molecular, development, and breeding disorders in zebrafish. Table 2 summarizes the articles reviewed in this section.

**Table 2.**
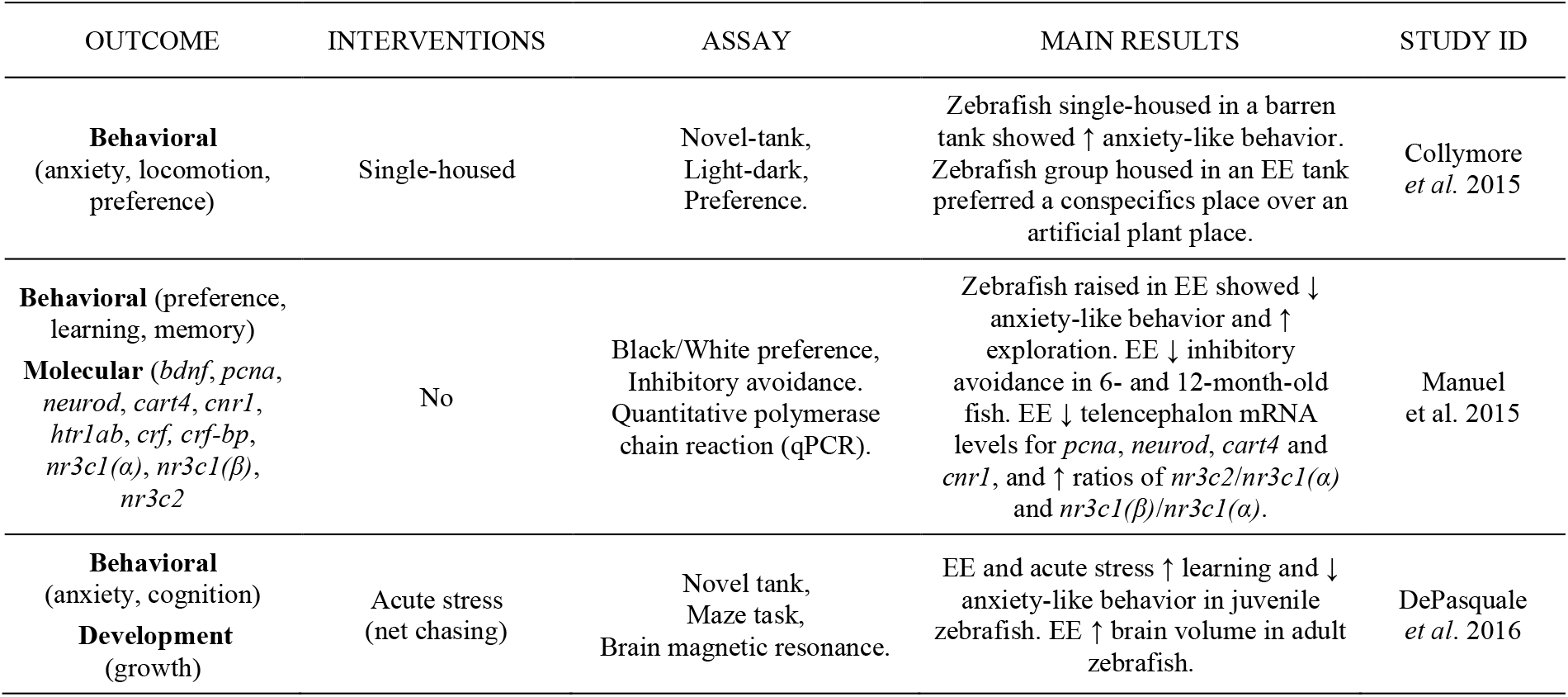
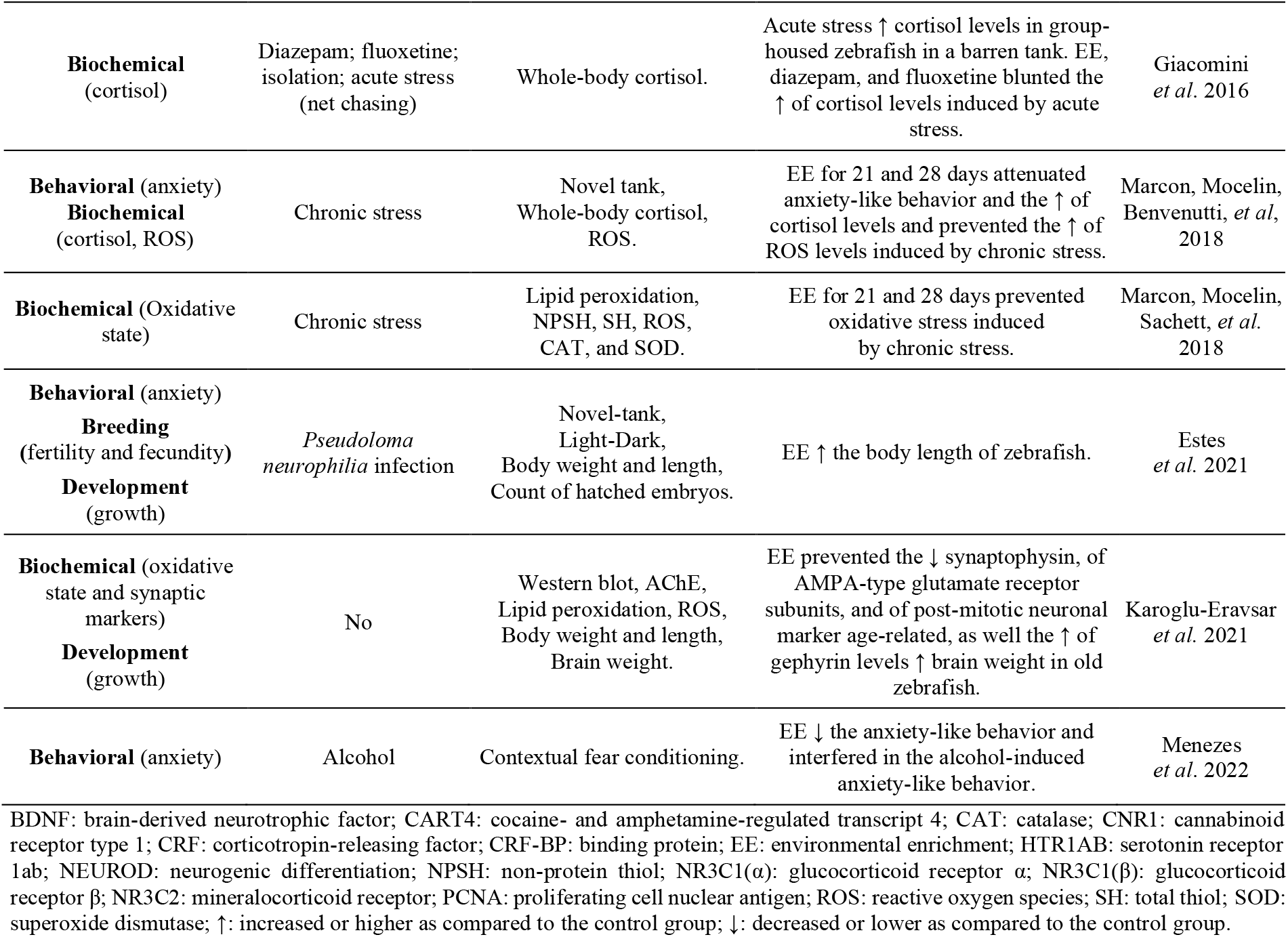
Studies included in this review that evaluated the EE effects on the prevention of disorders.

Stressful interventions such as isolation (Collymore et al., 2015), acute (DePasquale et al., 2016), and chronic stress (Marcon, Mocelin, Benvenutti, et al., 2018) increased anxiety-like behaviors in zebrafish, while EE exposition prevented these behavioral changes (Collymore et al., 2015; DePasquale et al., 2016; Marcon, Mocelin, Benvenutti, et al., 2018; Menezes et al., 2022). A useful tool to assess anxiety-like phenotypes in zebrafish is the light-dark test. This test is based on the scototaxis and the approach-avoidance conflict (Maximino et al., 2011). In general, when exposed to the light/dark apparatus, the adult zebrafish presents a preference for the dark compartment, while avoiding the light compartment (Maximino et al., 2011). Interestingly, anxiolytic drugs increase the time that zebrafish spend in the light compartment while anxiogenic drugs decrease this time (Sackerman et al., 2010). Manuel and collaborators (Manuel et al., 2015) found that zebrafish raised in an EE tank since hatching spent more time in the light compartment in the light-dark test when compared with zebrafish raised in a barren tank since hatching, which suggests that EE decreased anxiety-like behavior. Menezes and collaborators reported that EE exposure for 15 days prevented the anxiety-like behavior shock-induced in adult zebrafish, decreasing the time of freezing when compared to the barren tank exposure zebrafish for the same period. They also showed that EE exposure for 15 days prevented the freezing and locomotor behavior alterations alcohol-induced in adult zebrafish (Menezes et al., 2022).

Manuel and collaborators (Manuel et al., 2015) also demonstrated that 6 months post-fertilization (mpf) and 12 mpf zebrafish raised in an EE tank since hatching showed better learning and memory performance in an inhibitory avoidance test than those raised in a barren tank since hatching. While memory and learning impairment in aging is a normal physiological process, this data suggests that EE prevented the cognitive impairment related to the age advancement in 6 mpf and 12 mpf zebrafish. On the other hand, no differences were observed in 24 mpf zebrafish raised in an EE or a barren tank. The cognitive improvement EE-induced observed by Manuel and collaborators agrees with DePasquale and collaborators (DePasquale et al., 2016). They showed that juvenile zebrafish 25 days post-fertilization (dpf) exposed to EE for 60 days presented improvement of learning and memory performance when compared to juvenile zebrafish the same age exposed to a barren tank. These cognition parameters were evaluated in a maze task after zebrafish from both groups were exposed to acute stress by chasing (DePasquale et al., 2016).

While acute stress (net chasing for 120s) increased the whole-body cortisol levels in zebrafish housed in a barren tank, the same did not happen in zebrafish previously housed in an EE tank for 15 days, suggesting that EE blunted stress response (Giacomini et al., 2016). This result agrees with other reports showing that EE during 21 and 28 days attenuated the increase in the whole-body cortisol levels in zebrafish induced by an unpredictable chronic stress protocol (UCS) (Marcon, Mocelin, Benvenutti, et al., 2018). Apart from the prevention of hormonal changes, EE was also shown to prevent morphological alterations such as in the body length of zebrafish exposed to pathogens (Estes et al., 2021).

Marcon and collaborators (Marcon, Mocelin, Sachett, et al., 2018) showed that adult zebrafish subjected to a UCS protocol had an increase in lipid peroxidation and reactive oxygen species (ROS) levels and a decrease in non-protein sulfhydryl groups (NPSH) levels and superoxide dismutase (SOD) activity. These neurochemistry alterations are related to oxidative damage, and, interestingly, zebrafish pré-exposition to EE during 21 and 28 days prevented the decrease in NPSH levels and the increase in lipid peroxidation and ROS levels induced by UCS. Furthermore, pre-exposition to EE for 28 days also prevented the decrease in SOD activity UCS-induced, while both pre-expositions to EE for 21 and 28 days increased catalase (CAT) activity (Marcon, Mocelin, Sachett, et al., 2018). These data show that at the molecular level, improving housing conditions with EE is also able to induce deep effects on oxidative status, preventing oxidative damage induced by stress. Manuel and collaborators (Manuel et al., 2015) also showed that complex environments modulated the expression of different genes in 6-month-old zebrafish, leading to a reduction in the expression of *pcna* and other neuroplasticity-related genes such as neurogenic differentiation (*neurod*), cocaine- and amphetamine-regulated transcript 4 (*cart4*), and cannabinoid receptor 1 (*cnr1*) while an increase in the ratios of mineralocorticoid receptor (*nr3c2*)/glucocorticoid receptor *α* [*nr3c1(*α*)*] and glucocorticoid receptor *β* [*nr3c1(*β**)]/glucocorticoid receptor *α* [*nr3c1(*α*)*] was seen for fish under enrichment conditions.

Karoglu-Eravsar and collaborators (Karoglu-Eravsar et al., 2021) evaluated adult (6 mpf) and old (27 mpf) zebrafish exposed to EE for 4 weeks. They described global changes in cellular and synaptic markers age-related in zebrafish, such as to decrease in synaptophysin, in the AMPA-type glutamate receptor subunits, and in the postmitotic neuronal marker, and showed that the short-term pre-exposition to EE prevented these changes in old zebrafish. They also found that EE exposition increased the gephyrin level in old zebrafish. Gephyrin is an inhibitory scaffolding protein that plays a role as a structural and functional component in inhibitory synapses, and its increase may be related to a protective role against oxidative stress (Groeneweg et al., 2018). In addition, they showed that EE exposure increased brain weight in an age-dependent manner (Karoglu-Eravsar et al., 2021) Old zebrafish pré-exposure to EE showed an increase in brain weight when compared to those in the same condition without EE pré-exposure. However, the same was not observed in adult zebrafish (Groeneweg et al., 2018). In contrast, another study demonstrated that adult zebrafish (6 mpf) raised in EE from hatching showed an increase in brain volume compared to those raised in a barren tank (DePasquale et al., 2016). These results suggest that the effect of EE may depend on the zebrafish’s age and/or the duration of EE exposure. It is known that the first 15 days post-fertilization of zebrafish are essential for normal development, and EE exposure during this period seems to have a more robust effect in preventing further harmful effects.

In general, the results showed that EE was able to prevent some dysfunctions related to stressful stimuli, such as social isolation and acute and chronic stress protocols, alcohol exposure, conflict anxiety test, memory and learning impairment, and aging in pair-housed zebrafish. Therefore, these data suggest that EE intervention should be considered to prevent a series of behavior, physiological, and neurochemistry disorders in zebrafish.

### 3.3 EE protocols in zebrafish

With the popularization of the EE implementation, more and more researchers are applying this strategy to better understand its effects on laboratory animals either as a welfare improvement strategy or as a prevention strategy of behavioral, biochemical, molecular, and morphological alterations. Of the 27 studies included in this review, it is possible to notice that there is significant heterogeneity of protocols in studies with EE and zebrafish (table 3). The main variabilities found in these protocols are related to the materials used, the use or no of rotation of EE objects, exposure time, and the dimensions of the EE tank.

**Table 3.** EE protocols used in the studies included in this review.

| ENRICHMENT MATERIALS | EE<br>ROTATION | EXPOSURE<br>TIME | TANK<br>(L × W × H cm) | STUDY ID |
| --- | --- | --- | --- | --- |
| Artificial plant; English sea stones. | Unclear | 7 days | 15 L | von Krogh<br>*et al.* 2010 |
| Clay pots; natural plant<br>(*Ceratopteris thalictroides*); sand. | Unclear | 8 days | 130 × 50 × 50<br>100 × 50 × 50<br>160 × 41 × 50 | Kistler<br>*et al.* 2011 |
| Artificial plant. | Unclear | 6 months | 40 × 30 × 25 | Spence *et al.* 2011 |
| Artificial plant; gravel; gravel image; stone; sand; sand image. | Unclear | 7 days | 26 × 22 × 14,5 | Schroeder *et al.* 2014 |
| Artificial plant. | Unclear | 5 weeks | Unclear | Collymore *et al.* 2015 |
| Plastic vegetation. | Unclear | 1, 5, or 10 days | 6 × 11 × 16 | Keck *et al.* 2015 |
| Artificial plant; artificial stone; sand. | No | 12 months | 2 L<br>50 × 50 × 60 | Manuel *et al.* 2015 |
| Artificial plant; gravel; plastic bottle; plastic shelter; PVC pipe; stone. | Yes | 78 days | 35 x 19 x 28 | DePasquale *et al.* 2016 |
| Caps for refuge; gravel; natural plant; sand. | Unclear | 15 days | Unclear | Giacomini *et al.* 2016 |
| Artificial plant; plastic grass. | Unclear | Overnight | 1.5 L | Wafer *et al.* 2016 |
| Artificial plant; gravel; ruin-like plastic object. | No | 21 and 28 days | 40 × 20 × 24 | Marcon, Mocelin, Benvenuti, *et al.* 2018 |
| Artificial plant; gravel; ruin-like plastic object. | No | 21 and 28 days | 40 × 20 × 24 | Marcon, Mocelin, Sachett, *et al.* 2018 |
| Artificial plant; half clay pot; PVC pipe. | Unclear | 14-17 days | 20.8 L | Sykes *et al.* 2018 |
| Artificial plant; gravel; sand. | Yes | 6 weeks | 35 × 19 × 28 | DePasquale *et al.* 2019 |
| Artificial plant; gravel; natural plants (*Hypnaceae* sp., *Taxiphyllum Barbieri*). | Unclear | During testing | 100 × 25 × 25 | Jones *et al.* 2019 |
| Gravel; natural plants (*Vallisneria* spp., *Cryptocoryne wendtii*). | Unclear | 126 dpf | 33,5 × 19,5 × 17<br>21 × 13 × 13 | Lee *et al.* 2019 |
| Artificial plant; plant pot; sea image. | Unclear | 9 weeks | 30 × 15 × 24 | Woodward *et al.* 2019 |
| Artificial plant; PVC pipe; stone. | Unclear | 7 and 14 days | 40 × 25 × 16 | dos Santos *et al.* 2020 |
| Artificial plant; dark paper; marbles; marbles image; mirrored paper; PVC pipe; plants image; white tulle; zebrafish image. | Unclear | 7 days | 3 L | Krueger *et al.* 2020 |
| Artificial plant; PVC pipe. | Unclear | 3 days | 48 × 25 × 24 | Tan *et al.* 2020 |
| Artificial plant; grave; shelter. | Yes | 6 months | 35 × 19 × 28 | DePasquale *et al.* 2021 |
| Artificial plant. | Unclear | 5 weeks | 1.1 L | Estes *et al.* 2021 |
| Artificial plant; gravel; sea image; swimming tubes. | Unclear | 4 weeks | 50 x 15 x 30 | Karoglu-Eravsar *et al.* 2021 |
| Lego® (multiple colors and shapes). | Yes | 7, 14, and 21 days | 14 × 7 × 4 | Gatto, Dadda, *et al.* 2022 |
| Lego® (multiple colors and shapes). | Yes | 14 days | 14 × 7 × 4 | Gatto, Bruzzone, *et al.* 2022 |
| Artificial plants (orange, red, green); Background colors (red, yellow, blue, or green). | No | 10 days | 27 cm diameter | Maia *et al.* 2022 |
| Artificial plants; Lego®; marbles; PVC pipe. | Unclear | 15 days | 50 × 30 × 30 | Menezes *et al.* 2022 |

DPF: days post-fertilization; EE: environmental enrichment; H: height; L: length; W: width.

Figure 4 shows the great variety of enrichment materials that compose the protocols. Artificial plants are the most used artifacts to introduce complexity to the environment. As dense vegetation is found along the water streams where zebrafish are naturally found (Parichy, 2015) it is logical to perceive the importance of this material and why artificial plants are the most common items, being present in 21 protocols (77.78%). Natural plants are only used in 4 protocols (14.81%), as live plants can modify water quality. Other materials include gravel, which is used in 10 studies (37.04%), PVC pipes in 6 (22.22%), sand in 5 (18.52%), and printed images in 4 (14.81%). Lesser used items include Legos®, marbles, stones, and ruin-like plastic objects. An important strategy derived from this topic must be considered. Not only the types of materials used for the enrichment are important but so is their presentation to the animals (Roy et al., 2022). Rotation of EE can increase the attraction of the fish to the items presented, elevating the complexity of the environment, and diminishing the habituation effect to the environmental conditions. This strategy was only implemented in 5 studies (18.52%) and thus should be better explored in future research.

**Figure 4.**
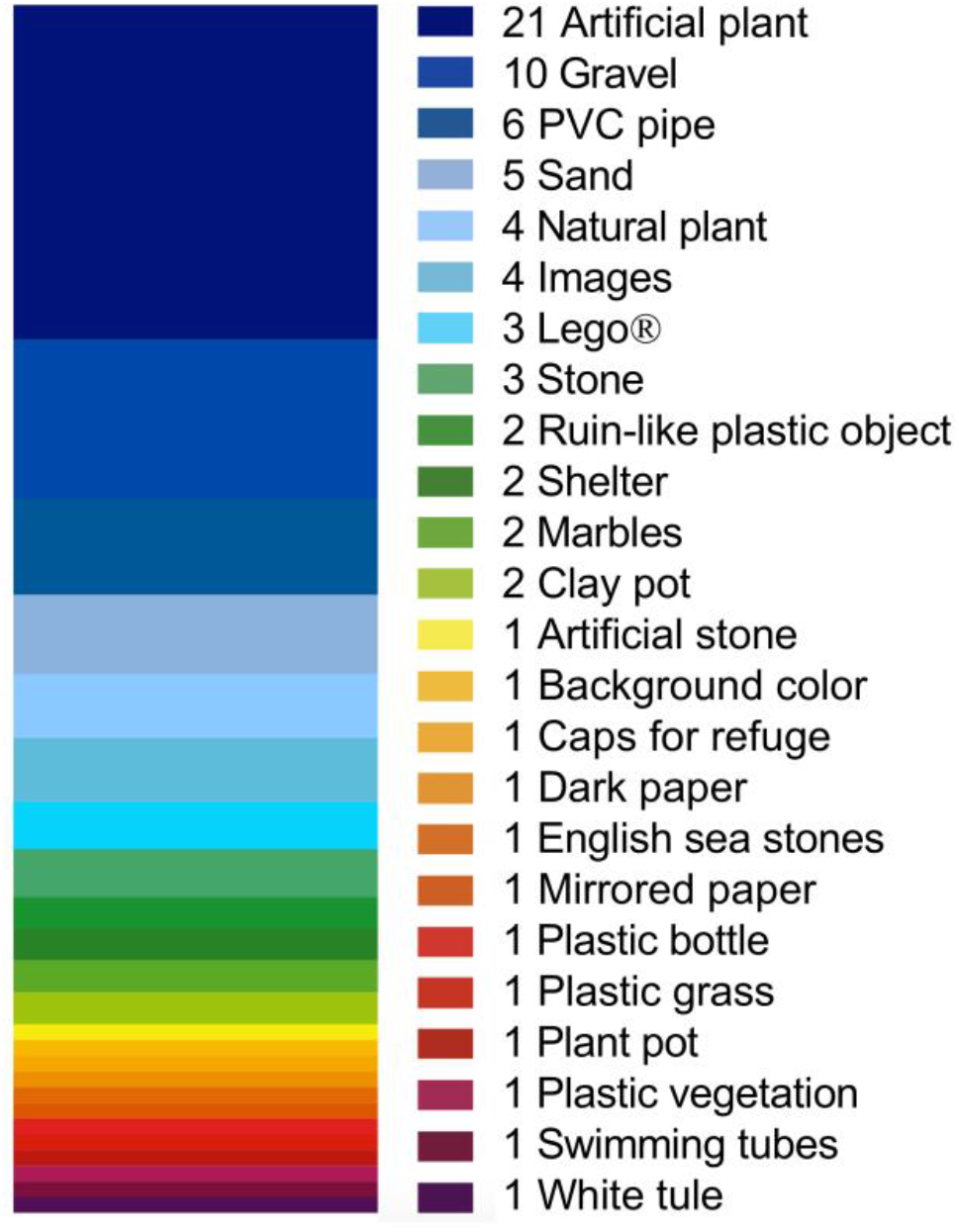
Different materials were used to compose the EE in zebrafish studies for any results published in peer-reviewed scientific journals between 2010 and 2021. The number of studies that used each type of material is found before each material.

The exposure time of the animals to the enriched habitat varied widely, with 9 studies involving experiments with fish exposed to complex housing conditions for up to a week (33.33%), and 21 studies (77.78%) for a longer period. In 6 studies (22.22%) more than one exposure duration regimen was used. Interestingly, the shortest protocol exposed fish to an EE aquarium only during the outcome assessment to evaluate the preference between enrichment conditions (Jones et al., 2019). The longest protocol exposed zebrafish from hatching up until the animals completed one year of life (Manuel et al., 2015). Independently of the duration, zebrafish showed a significant preference for complex environments (DePasquale et al., 2019; Tan et al., 2020) and EE prevented several deleterious alterations (DePasquale et al., 2016; Manuel et al., 2015). Apart from protocol duration, many differences are also found in the literature for the shapes and dimensions of housing tanks used in these experiments.

The diversity found in the experimental conditions that fish are exposed to in the included studies is also summarized in table 4. Habituation to lab settings is used to avoid acute effects of transportation to and housing in a novel environment in the behavior of animals. Between the studies selected, habituation time varied from zero days to 5 months. Since reproduction is often carried out in the same facilities where outcome assessment is conducted and animals are sometimes introduced to the EE as soon as hatching takes place, 9 studies (33.33%) do not report habituation time. Another common scheme is to use 14 to 15 days of habituation, as used in 6 experiments (22.22%). The zebrafish diet is a more homogenous topic with common practices involving feeding the animals with flaked food accompanied by the brine shrimp *Artemia salina* as in 17 studies (62.96%), which covers the standard diet described in specialized literature (Westerfield, 2007). Feeding frequency can vary from one to four times a day.

**Table 4.**
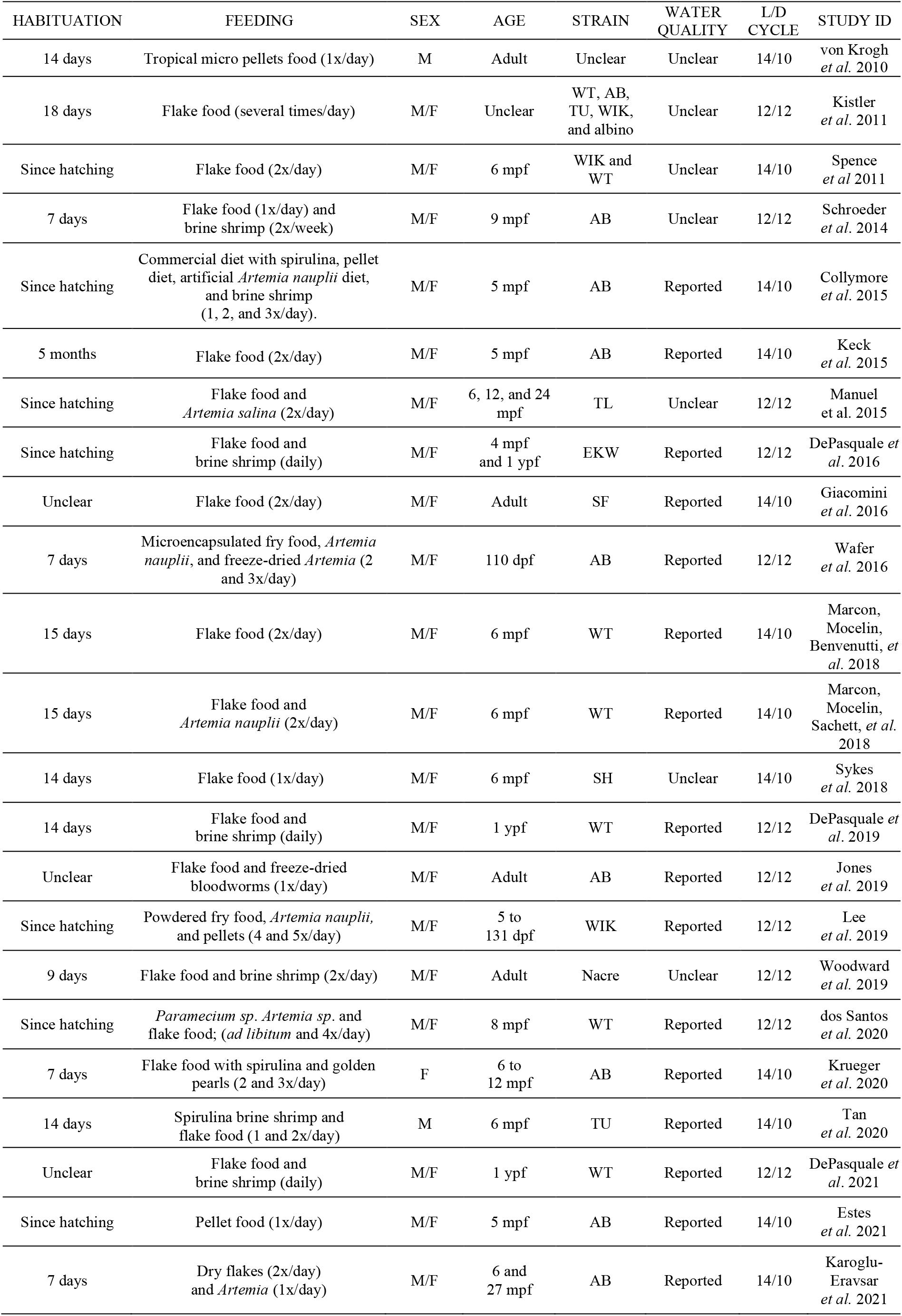
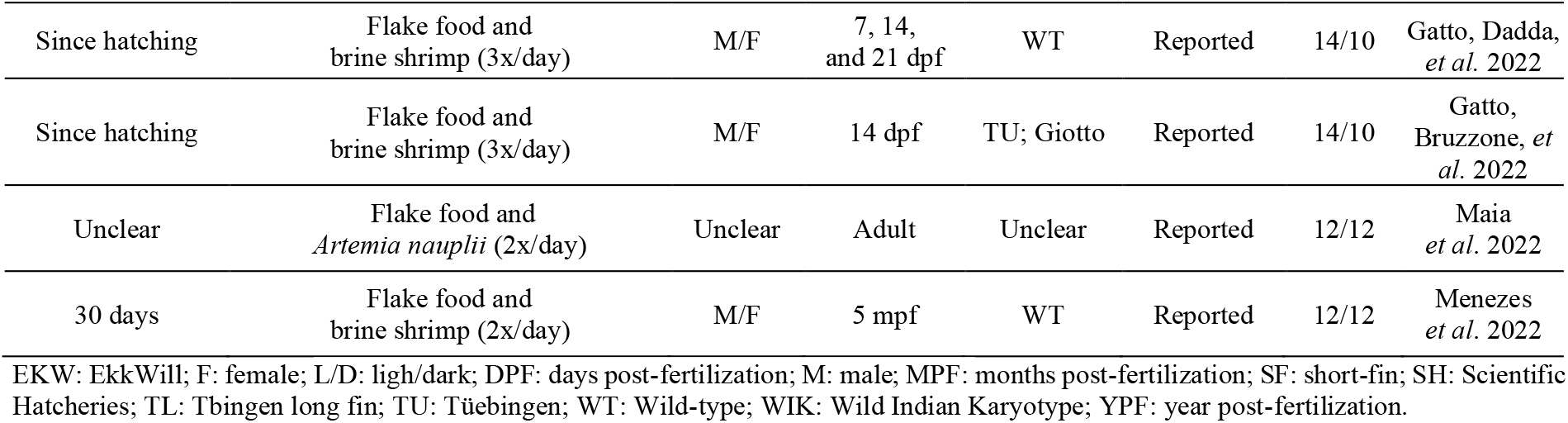
Experimental details of the studies that were included in this review.

Researchers used a mix of male and female zebrafish in 23 studies (85.18%), which resembles population structure in natural conditions and avoids sex-specific biases from housing animals of only a specific sex together. Only male fish were used in 2 experiments (7.41%), while only females were used in 1 (3.70%). Outcome assessment was carried out using exclusively adults in 23 of the experiments reported (85.18%). This points out the importance of better exploring the EE effects directly in early life stages. The most common strains of zebrafish used in the experiments were Wild-type and AB with both being used in 9 studies (33.33%). Some of the other strains used were WIK, TU, and EW.

Water quality is one of the most important parameters to be controlled when working with fish as a model animal. Lack of standardization in practices to ensure the optimal conditions for the organism have a huge impact on welfare and on the desired outcomes to be assessed in the experiments. Seven studies (25.93%) did not report water quality information, raising questions on the potential bias associated with the results. Lighting conditions were reported in all studies in the review and strategies were evenly split with 14 studies (51.85%) using a regimen of 14/10 hours light/dark cycle and 13 (48.15%) with 12/12 hours light/dark cycle. This is an additional criterion to be standardized in other to optimize the evaluation of desired effects without the interference of the light/dark cycle since the circadian cycle has a direct impact on physiology and behavior (Kopp et al., 2018; Krylov et al., 2021).

The substantial heterogeneity between protocols used in preclinical biomedical research is often pointed to as the source of reproducibility problems. This is no different for experiments using zebrafish since studies such as those included in this review frequently lead to conflicting results (Gerlai, 2019). This calls for an improvement in research techniques and the report of all procedures, lab conditions, apparatuses, and materials used in experiments (Benvenutti et al., 2021) as well as adhering to practices to reduce the risk of bias associated with the methodological conduct (Macleod et al., 2015).

### 3.4 Risk of bias

The overall risk of bias for the items evaluating the methodological quality of included studies (items 1 to 5) was considered low, except for outcome blinding and the baseline characteristics of the animals (Figure 5). Randomized allocation of animals to the experimental groups was reported in 21 studies (77.78%) while 6 studies (22.22%) did not provide sufficient information to rule out bias. For baseline characteristics, only 13 studies (48.15%) presented a low risk of bias. On the other hand, 14 studies (51.85%) were considered as presenting an unclear risk of bias due to the lack of information on the characteristics of included animals. Bias related to the blind assessment of outcomes was considered low in 9 studies (33.33%) and unclear in 18 studies (66.67%). It was unclear whether available data was incomplete in 5 studies (18.52%), but most studies (81.48%) scored a low risk of bias for this item. Similarly, for selective reporting, most studies (92.59%) scored a low risk of bias, showing that the results were consistent with the methodology reported.

**Figure 5.**
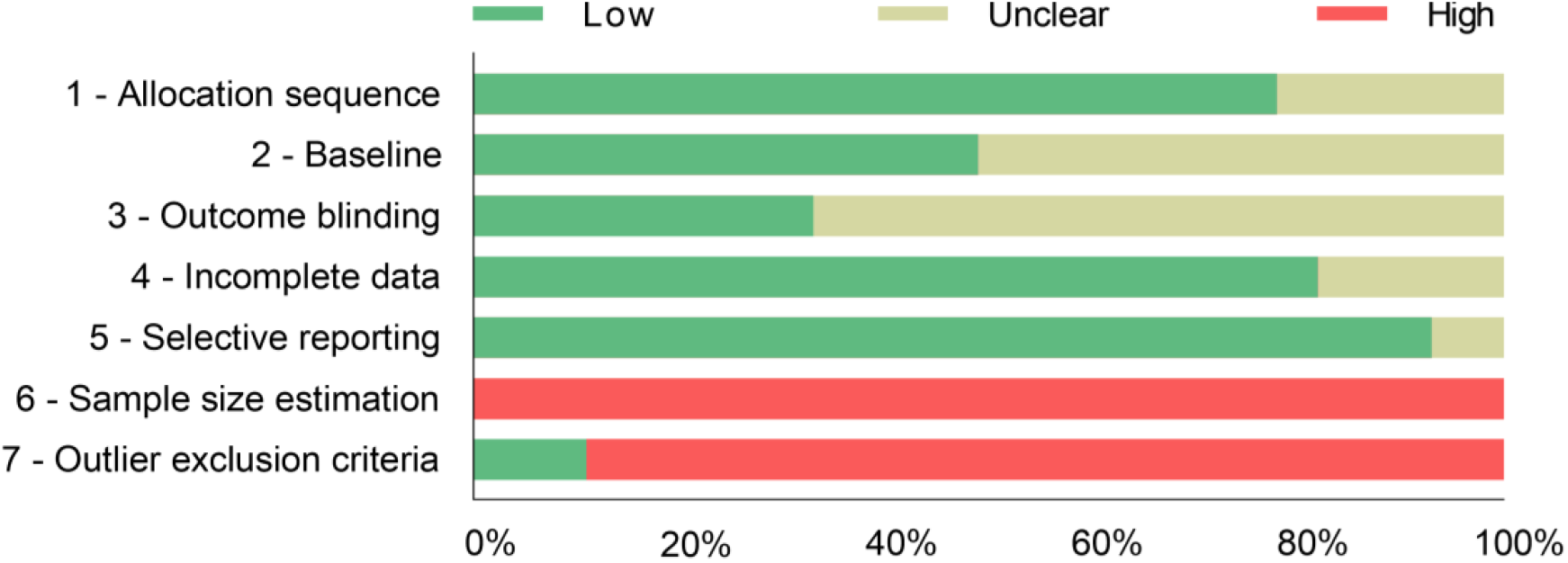
Risk of bias assessment of included studies. The risk of bias assessment was performed based on the SYRCLE’s risk of bias assessment tool. Items were scored as presenting a high, unclear, or low risk of bias. Classification is given as the percentage (%) of assessed studies (n = 27) presenting each score.

Contrasting with the methodological quality, the reporting quality of the included studies for the selected items was considered overall as presenting a high risk of bias (Figure 5). Although reporting the methodology implemented for sample size estimation in the studies and the existence or absence of any outlier exclusion criteria are important conducts while describing experiments, 27 studies (100%) did not inform the procedures for estimating the number of animals to be used in the experiments and 24 (88.89%) lacked the criteria for excluding animals from the analyses.

## 4 CURRENT CHALLENGES AND FUTURE RESEARCH

Therefore, we summarized some basic scientific questions that are still open and must be answered in future studies about the applicability of EE in zebrafish research.

### 4.1 Unanswered scientific questions related to EE and zebrafish

➔ What are the best EE materials to apply in zebrafish homes in a laboratory environment?
➔ How long should EE be applied to laboratory zebrafish?
➔ Is there a specific developmental stage where EE is most beneficial to zebrafish?
➔ Is it necessary to rotate EE materials? And how often?
➔ Will it be possible to apply EE to the zebrafish rack closed system?

As these scientific questions are being answered and new standardized experimental protocols emerge, we believe that this knowledge must be applied in the zebrafish facilities and laboratory tanks. According to the results described here, the use of EE in the zebrafish home tank provides better welfare and may reduce sources of bias in scientific experiments, such as high-stress levels and fighting events. In fact, due to animal welfare and its ethical and scientific benefit, the practice of adding enrichment to the home in the laboratory environment is already recommended by several regulatory agencies for the control of animal experimentation in the world, thus being used in several animal facilities and research centers, mainly for rodents.

## 5 CONCLUSION

Although the zebrafish EE protocols reviewed in this review presented a series of experimental differences, the results showed that the benefits of the EE for zebrafish were robust. However, the necessity to develop standardized protocols, aiming to minimize possible bias sources in scientific research has been increasingly discussed. Hence, it is important to emphasize that many questions in this field remain misunderstood. We believe that studies should go beyond the description of the results and should also describe in detail their methodology to facilitate the development of replicable and standardized protocols to better apply the EE in scientific studies with zebrafish.

